# Thyroid hormone coordinates developmental trajectories but does not underlie developmental truncation in danionins

**DOI:** 10.1101/562074

**Authors:** Yinan Hu, Angela Mauri, Joan Donahue, Rajendra Singh, Benjamin Acosta, Sarah McMenamin

## Abstract

**Background:** Differences in postembryonic developmental trajectories can profoundly alter adult phenotypes and life histories. Thyroid hormone (TH) regulates metamorphosis in many vertebrate taxa with multiphasic ecologies, and alterations to TH metabolism underlie notable cases of paedomorphosis in amphibians. We tested the requirement for TH in multiple postembryonic developmental processes in zebrafish, which has a monophasic ecology, and asked if TH production was compromised in paedomorphic *Danionella*.

**Results:** We showed that TH regulates allometric growth in juvenile zebrafish, and inhibits relative head growth. The lateral line system showed differential requirements for TH: the hormone promotes canal neuromast formation and inhibits neuromast numbers in the head, but causes expansion of the neuromast population in the trunk. While *Danionella* morphology resembled that of larval zebrafish, the two *Danionella* species analyzed were not similar to hypothyroid zebrafish in their shape or neuromast distribution, and both possessed functional thyroid follicles.

**Conclusions:** Although zebrafish do not undergo a discrete ecological transformation, we found that multiple tissues undergo transitions in developmental trajectories that are dependent on TH, suggesting the TH axis and its downstream pathways as likely targets for adaptation. Nonetheless, we found no evidence that evolutionary paedomorphosis in *Danionella* is the result of compromised TH production.

**Bullet Points:**

- Thyroid hormone regulates shifts in relative growth trajectories in different zebrafish tissues
- Thyroid hormone inhibits head growth in juvenile zebrafish, and regulates juvenile growth patterns
- Thyroid hormone stimulates formation of canal neuromasts in the head and proliferation of trunk neuromasts in zebrafish
- *Danionella*, a miniaturized danionin lineage closely related to zebrafish morphologically resemble larval zebrafish and do not form neuromast canals
- Hypothyroidism is not the cause of paedomorphosis in *Danionella*

**Grant Sponsors:** NIH R00GM105874

NIH R03HD091634

Burroughs Wellcome Collaborative Research Travel Grant 1017439 NSF CAREER 1845513

## INTRODUCTION

Shifts in the timing and magnitude of developmental processes during later, postembryonic stages of development can profoundly alter adult phenotypes (Gould, 1977; Raff, 1996; Smith, 2001). Revealing the pathways that regulate ontogenetic trajectories of different developmental processes will be critical in understanding how life histories and phenotypes are formed. Further, discerning the mechanisms by which different processes may be decoupled will be critical for a complete understanding of the evolution of development.

An extreme way in which postembryonic development may be changed is through paedomorphosis, wherein an organism does not complete terminal stages of development (Gould, 1977; Shaffer and Voss, 1996); organisms on the spectrum of paedomorphosis retain larval characteristics into adulthood (Johnson and Voss, 2013). Paedomorphosis is generally most obvious when it originates in clades with ecologically multiphasic life histories. Ecologically multiphasic vertebrates are those that undergo an ecological shift as they transform to juveniles, e.g. amphibians that metamorphose from aquatic larvae to terrestrial juveniles. Paedomorphosis in these clades involves loss of a markedly divergent life history stage and may be phenotypically quite obvious (McMenamin and Hadly, 2010; Johnson and Voss, 2013). Paedomorphosis in clades with ecologically monophasic life histories may be less morphologically striking, but paedomorphosis has nonetheless been posited as an important evolutionary process in many monophasic groups, including in mammals (e.g. Shea, 1983; Goodwin et al., 1997).

Since vertebrates with multiphasic life cycles often undergo a metamorphosis triggered by a single molecular pathway (generally thyroid hormone, TH), paedomorphosis could theoretically evolve in these groups through a single genetic change. Thus, disruptions to TH pathways cause developmental arrest and underlie cases of paedomorphosis, as in the Mexican axolotl (see Johnson and Voss, 2013; De Groef et al., 2018). In contrast, ecologically monophasic vertebrates generally have more subtle relationships with the molecular signals that regulate progression through post-embryonic development. TH functions as an important regulator of developmental processes in ecologically monophasic vertebrates (e.g. zebrafish and mammals), but disruptions to the thyroid axis do not result in whole-organism arrest (Brown, 1997; McMenamin et al., 2014; Buchholz, 2015). In ecologically monophasic groups, the pathways regulating postembryonic developmental progression are less clear, as are the mechanisms and pathways that might underlie evolutionary paedomorphosis.

Teleosts represent the largest and most speciose group of vertebrates, exhibiting a remarkable range of life history strategies (Finn and Kapoor, 2008; McMenamin and Parichy, 2013). Most marine fishes are ecologically multiphasic, and pelagic larvae undergo profound physiological metamorphosis to transform into benthic juveniles (McMenamin and Parichy, 2013). Metamorphosis is stimulated by TH in flat fish (Isorna et al., 2009; Xu et al., 2016; Campinho et al., 2018) and reef fish (Holzer et al., 2017), and likely in many other species (Holzer and Laudet, 2015). Like in amphibians, TH stimulates the abrupt transformation of discrete tissues across the body, causing dramatic changes in overall morphology, sensory systems and other organs.

In contrast, most freshwater fishes, including zebrafish (*Danio rerio*), are ecologically monophasic. These monophasic species show a more subtle and protracted transformations from larva to juvenile. Nonetheless, specific tissues (including skeleton, pigment pattern and the lateral line) do undergo major remodeling during postembryonic development (Parichy and Turner, 2003; Webb and Shirey, 2003; Parichy, 2006; Parichy et al., 2009). Many of these remodeling processes, including those in the skeleton and the pigment pattern, are dependent on TH (Shkil et al., 2010; Shkil et al., 2012; McMenamin et al., 2014; Keer et al., 2019). Whether in total these transformations and remodeling processes constitute a “metamorphosis” depends on how broadly the term is defined (e.g. see Laudet, 2011; McMenamin and Parichy, 2013). A clearer picture of normal developmental trajectories in zebrafish, as well as the relative dependencies on TH, can inform our concept of how normal postembryonic development is regulated and whether this species exhibits a metamorphosis homologous to that of marine species.

Many fish species have been described as paedomorphic, including ice fish (Albertson et al., 2010), gobies (Johnson and Brothers, 1993; Giovannotti et al., 2007) and blennioid fishes (Hastings, 2002). *Danionella* is a danionin relative of zebrafish, sister clade to the group containing zebrafish and *Danios, Devarios* and *Microdeverios* (McCluskey and Postlethwait, 2014). *Danionella* are miniaturized and developmentally arrested, resembling larval zebrafish in several ways: adult *Danionella* lack scales, barbels and many typically late-ossifying bones (Britz et al., 2009; Britz and Conway, 2016), and retain a larva-like pigment pattern and feeding kinematics (McMenamin et al., 2017; Galindo et al., 2018). *Danionella translucida* is emerging as a novel model for studying behavior, neurobiology and brain function (Penalva et al., 2018; Schulze et al., 2018), and a hybrid genome assembly was recently completed (Kadobianskyi et al., 2019). A better understanding of the developmental relationships between *Danionella* and zebrafish, along with information about the pathways underlying paedomorphosis in *Danionella*, will offer new insights about this emerging model.

In this study we focused on two trait systems that change over the course of postembryonic development. We first focused on body shape, which changes dramatically during postembryonic development of many species. Vertebrates with multiphasic life cycles exhibit considerable shape change (i.e. allometric growth) during their profound physiological metamorphosis, followed by more proportional (i.e. isometric) growth as juveniles and adults.

We next focused on development of the lateral line system. This flow-sensing sensory system consists of individual neuromasts distributed across the body in specific patterns (Webb and Shirey, 2003; Coombs et al., 2014; Webb, 2014). The lateral line system is transformed during metamorphosis in amphibians (Wahnschaffe et al., 1987) and lampreys (Gelman et al., 2008), and in teleosts with both ecologically multiphasic (e.g. flatfishes; Harvey et al., 1992) and ecologically monophasic lifecycles (e.g. zebrafish; Webb and Shirey, 2003).

Our goals in this study were three-fold. <u>First</u>, we aimed to characterize shape change and the development of the lateral line system during normal zebrafish development, asking whether these developmental trajectories show evidence of a metamorphic transformation that corresponds with a period of increased TH production. <u>Second</u>, we tested whether TH is required in the development of these systems, determining the extent to which the hormone is required to coordinate shape change and/or lateral line development with overall growth. <u>Finally</u>, we tested whether adults of two *Danionella* species recapitulate any stage of normal zebrafish ontogeny, further testing the possibility that developmental arrest in *Danionella* is due to a lack of TH production.

## RESULTS AND DISCUSSION

### Zebrafish show allometric shifts during postembryonic development

To test the extent to which euthyroid wild-type (WT) zebrafish undergo changes in overall morphology during postembryonic development (see Fig 1A-B), we first assessed allometric growth trajectories of body shape (Fig 2, green crosses). Together, the first three principal components captured more than 88% of total shape variation; these were plotted against individual standard length (SL; Fig 2). PC1 primarily captured body depth (Fig 2D, G). In this axis, WT zebrafish allometrically increase body depth until the end of the larval period (10 mm SL), then maintain a roughly isometric change in body depth (Fig 2A). PC2 captures the relative size of the head. This component peaks at ~9 mm SL, and then decreases gradually during juvenile and adult growth (green crosses, Fig 2B). The dorsal profile, captured by PC3 increases gradually with growth (Fig 2 C).

**Figure 1.**
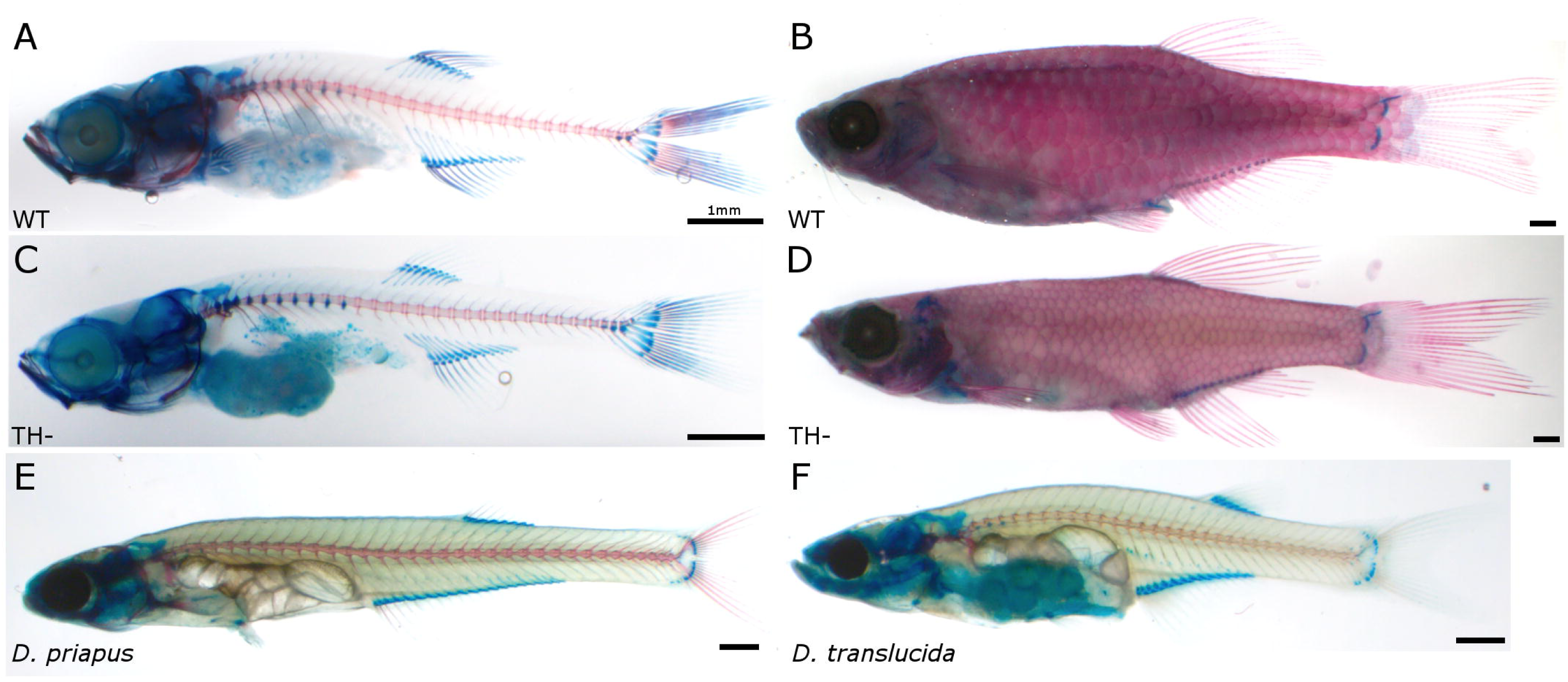
Cleared and stained zebrafish and *Danionella*. **A**, wild-type (WT) juvenile zebrafish. **B**, WT adult zebrafish. **C**, TH-juvenile zebrafish. **D**, TH-adult zebrafish. **E**, adult *D. priapus*. **F**, adult *D. translucida*.

**Figure 2.**
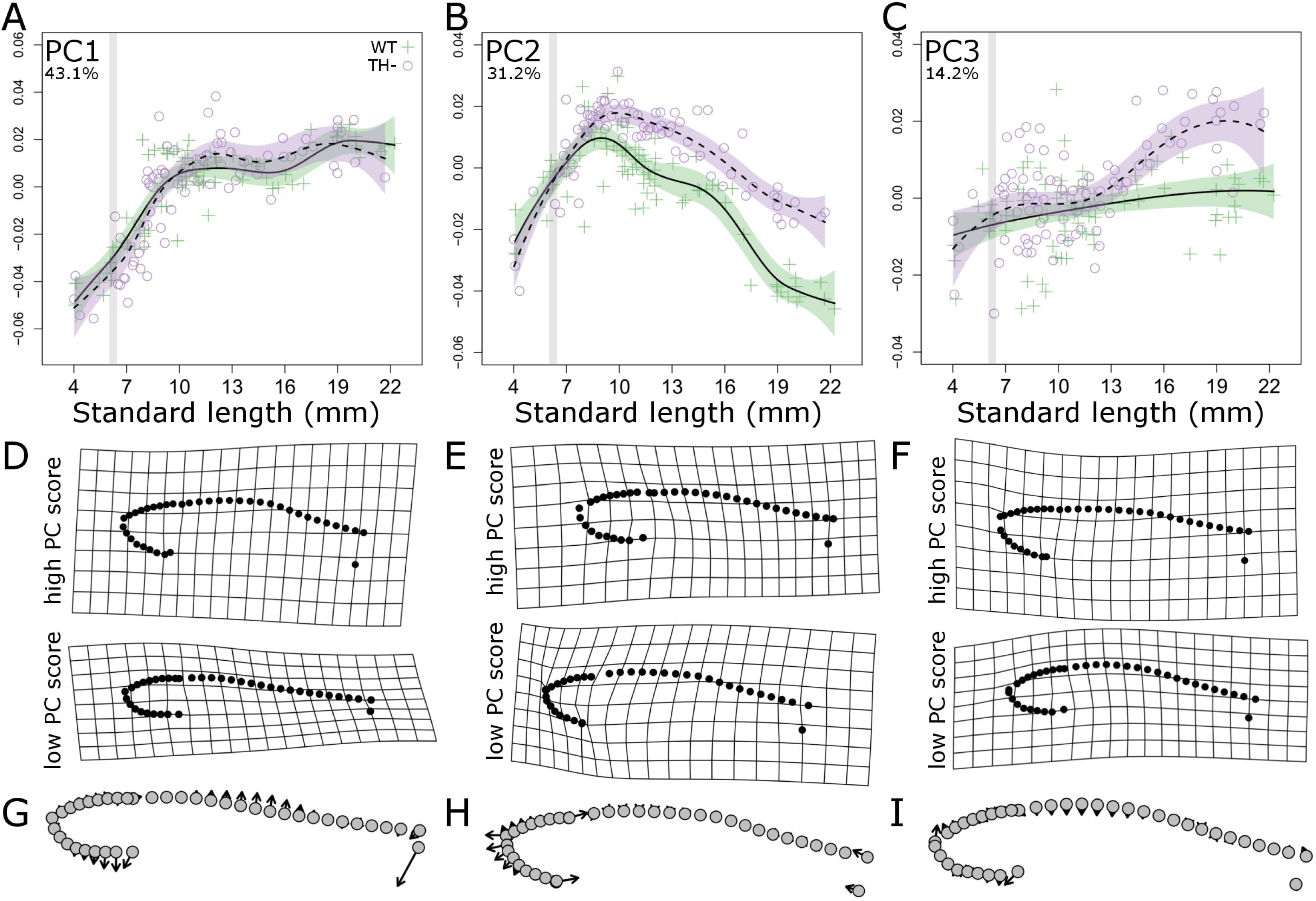
Thyroid hormone regulates allometric trajectories in zebrafish. **A-C**, Principle component (PC) scores of body shape plotted against standard length for both WT (green cross) and TH-(purple open circle) zebrafish. Percent variation explained by each PC axis is shown in the upper left corner. Developmental trajectory is shown in solid line with purple standard error for WT and dashed line with green standard error for TH-. For comparison between graphs, grey stripes highlight period from 6-6.4 SL, during which TH availability is elevated. **D-F**, deformation grids showing extreme shapes along each PC. **G-I**, vector plots showing shape change along each PC axis, with arrows pointing from negative to positive PC values.

To more directly test for growth in head proportion during development, we plotted the ratio of head length/SL over the course of growth (Fig 3C, green crosses). This ratio mirrored PC2, peaking at ~8 mm SL and then gradually declined afterwards (green crosses, Fig 3C). Eye size had a similar developmental trajectory to head length, continued to grow proportionally during larval phase, peaking at ~10 mm SL (green crosses, Fig 3D). Thus, WT zebrafish exhibit shifts in allometric relationships during the transition from larva to juvenile (8-10 mm SL); these shifts are consistent with a systemic metamorphosis. The pattern of positive allometric growth of the head has been documented in a number of fish species (e.g. see Osse et al., 1997; Gisbert, 1999; Gisbert and Doroshov, 2006), and likely reflects strong selection on early development of craniofacial feeding apparatus, respiratory and sensory systems.

**Figure 3.**
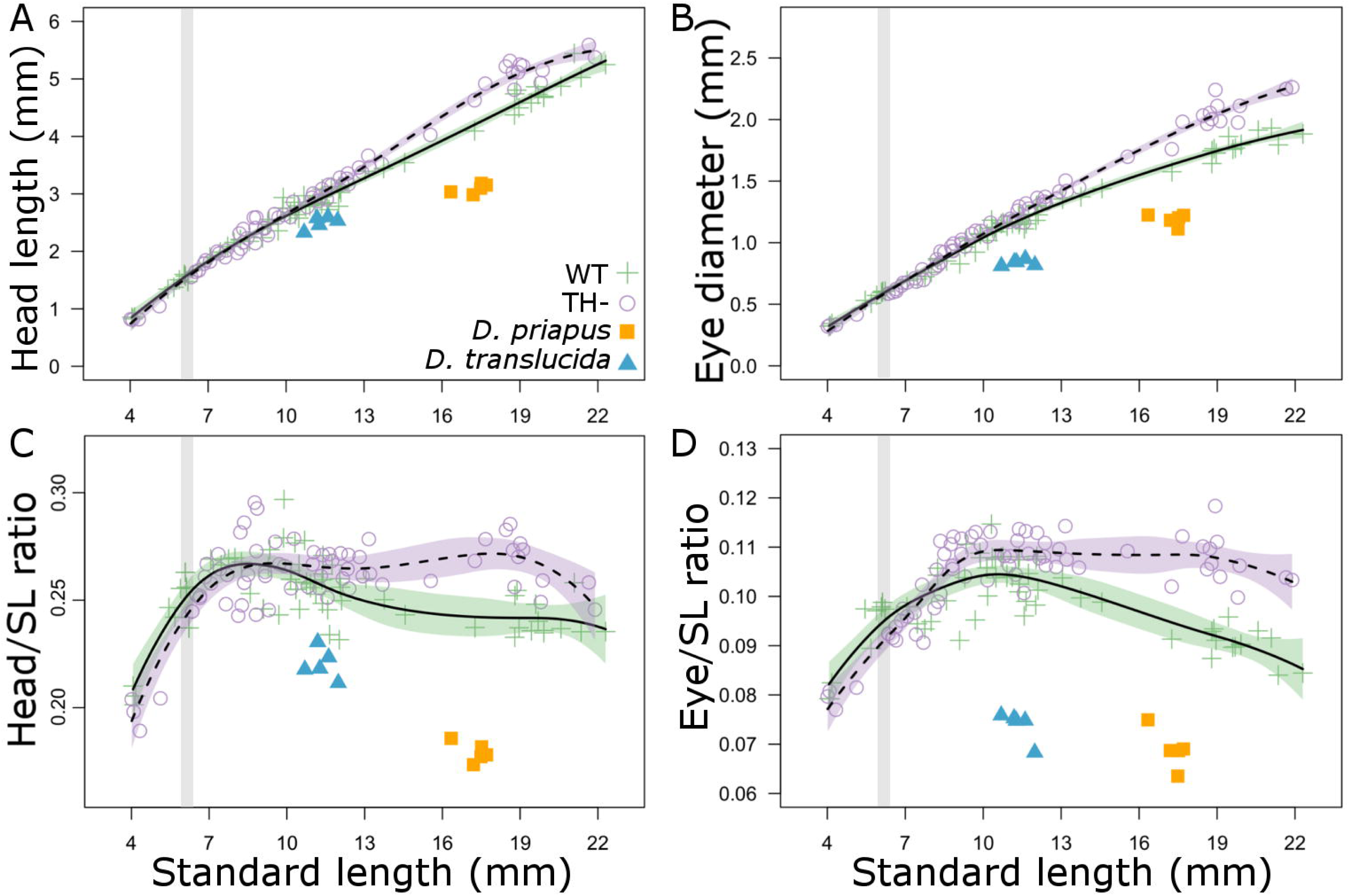
Thyroid hormone regulates body proportions in zebrafish. **A**, head length as a function of SL; **B**, eye diameter as a function of SL. **C**, Head length/SL and **D**, eye diameter/SL as functions of SL. Showing both WT (green cross) and TH-(purple open circle) zebrafish, with comparison to *D. priapus* (orange square) and *D. translucida* (blue triangle) adults. Developmental trajectory is shown in solid line with purple standard error for WT zebrafish and dashed line with green standard error for TH-zebrafish. For comparison between graphs, grey stripes highlight period from 6-6.4 SL, during which TH availability is elevated.

### TH inhibits head growth in the juvenile

To determine the extent to which TH is required for these allometric shifts, we measured shape and proportion changes in transgenically thyroid ablated, hypothyroid fish (TH-; Fig 1C, D). Changes in body depth were independent of TH, and shape changes along the PC1 axis were indistinguishable between TH- and WT (Fig 2A). However, we found that TH influences the relative growth of the head: in TH-fish, PC2 continues to increase along a larva-like trajectory until ~10 mm SL (purple circles, Fig 2 B). The third PC axis reveals that TH- fish develop an increasingly flat dorsal profile, unlike that of any stage of development in WT (Fig 2C).

In TH-fish, heads and eyes continue to grow along relatively steeper (larva-like) trajectories (Fig 2A, B). After rapid relative growth during larval development, the relative size of the head and eye does not decline appropriately in TH- fish (Fig 2C-D). Thus, TH is required for the coordinated transition from a larval trajectory to the juvenile trajectory, and limits the relative growth of the head in juveniles.

### TH inhibits proliferation of cranial neuromasts, but promotes formation of neuromast stitches on the body

We next focused on the development of a trait that emerges via very different processes than allometric body growth. We tracked numbers of neuromasts of the lateral line system across a range of body sizes (see Fig 4). In WT zebrafish, the number of trunk neuromasts increases roughly in proportion to body size, whereas neuromasts in the head show a negative allometric relationship to body size (green crosses, Fig 5A, B). We found that TH- adults had ~25% more total neuromasts than did WT adults (Fig 5C), and that this difference was driven primarily by the increasing number of cranial neuromasts (Fig 5A). This increase in cranial neuromast number is not simply due to the fact that TH- fish have larger heads, as differences remain even when standardized by head length (Fig 5D).

**Figure 4.**
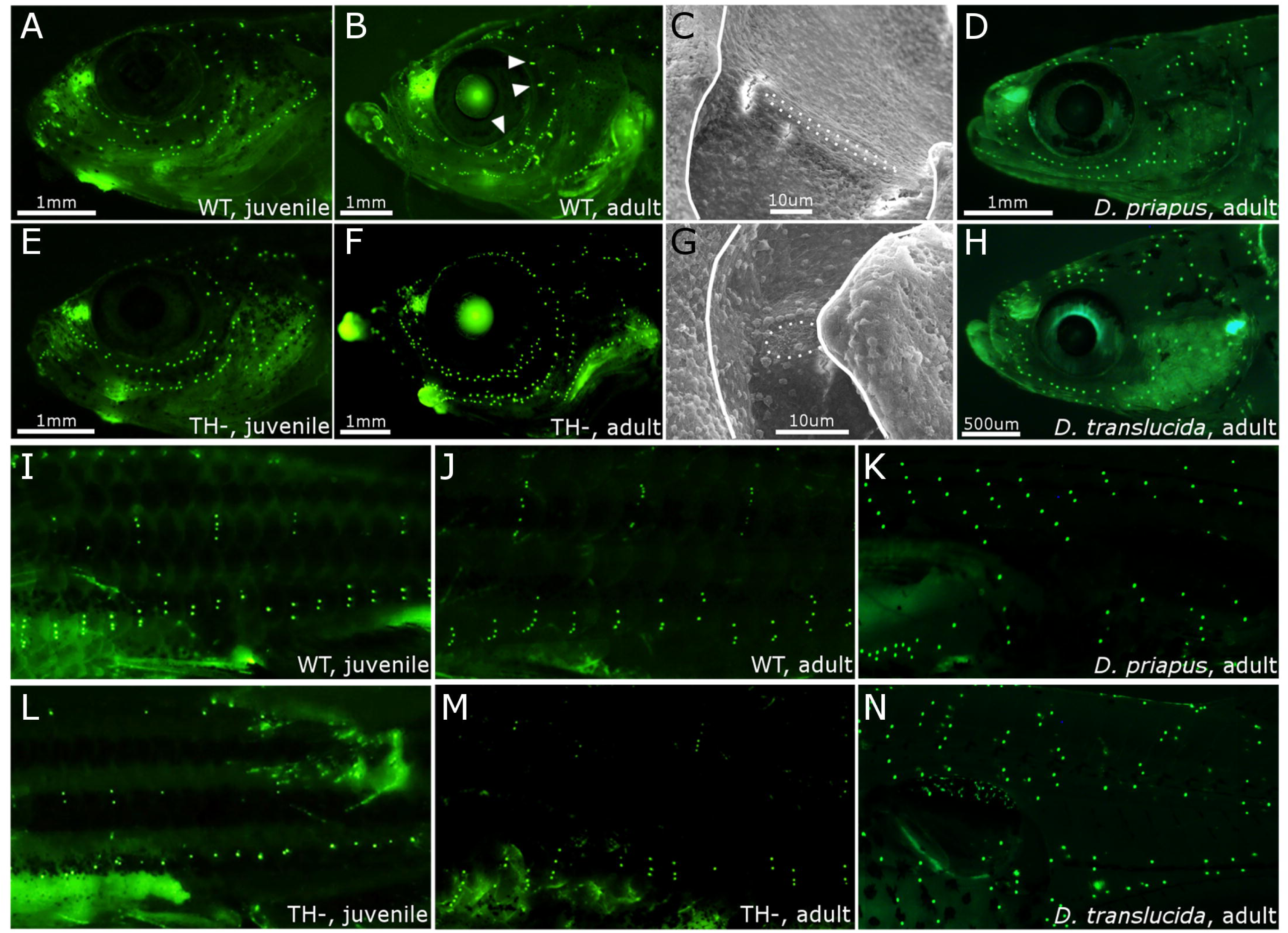
The lateral line system in zebrafish and *Danionella*. **A-B**, DiASP live staining of head neuromasts in WT juvenile (13 mm SL) and adult (18+ mm SL). Arrowhead indicates rod-shaped canal neuromasts at adult stage. **C**, Scanning electron micrographs of a supraorbital canal neuromast in WT adult. Dotted line outlines the neuromast; solid line outlines the bony edge of the canal. **D**, head neuromasts in *D. priapus* adult. **E-G**, same as B-D in TH-zebrafish. **H**, facial neuromasts in *D. translucida* adult. **I-N**, DiASP live staining of trunk neuromasts in WT juvenile zebrafish (13 mm SL, **I**), WT adult (18+ mm SL, **J**), *D. priapus* adult (**K**), TH-juvenile (13 mm SL, **L**) TH-adult (18+ mm SL, **M**), and *D. translucida* adult (**N**).

**Figure 5.**
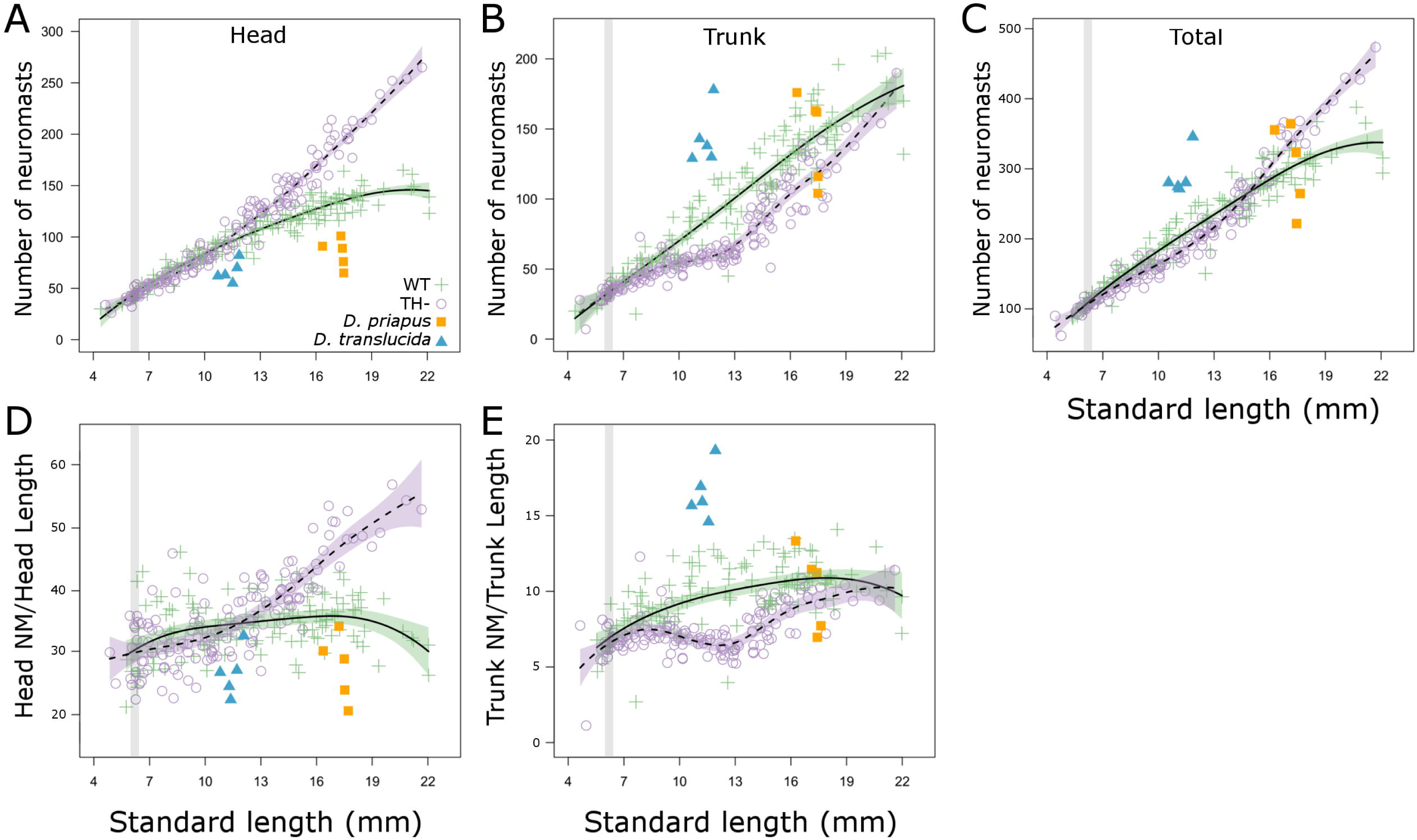
Thyroid hormone regulates lateral line development in zebrafish. **A**, number of cranial neuromasts, as a function of SL, with comparison to *D. priapus* (orange square) and *D. translucida* (blue triangle) adults. For zebrafish, developmental trajectory is shown in solid line for WT and dashed line for TH-. Developmental trajectory is shown in solid line with purple standard error for WT zebrafish and dashed line with green standard error for TH-zebrafish. For comparison between graphs, grey stripes highlight period from 6-6.4 SL, during which TH availability is elevated. **B-E**, same as A for number of trunk neuromasts, total number of neuromasts (head + trunk), cranial neuromasts divided by head length, and trunk neuromasts divided by trunk length.

A subset of cranial neuromasts acquire an elongated, rod-like shape in WT juveniles and eventually sink into bony canals (Webb and Shirey, 2003). We found that although neuromasts do become enclosed in canals in TH- adults, these canal neuromasts fail to acquire the mature morphology seen in WT (Fig 4B-C versus 4F-G). The transition into mature canal neuromasts may serve as a cue that inhibits neuromast proliferation; without this cue, neuromasts in TH-heads may remain highly proliferative through adulthood.

In WT zebrafish, trunk neuromasts proliferate into vertical stitches starting at ~10 mm SL (Ledent, 2002; visible in Fig 4I-J and M). In TH- fish, the trunk neuromast population grows more slowly until stitches start forming at ~13 mm SL (see Fig 4M and Fig 5B), suggesting that TH regulates onset of stitch formation.

### Bioavailability of TH shifts during mid-larval development

TH production peaks during metamorphic climax in amphibians, measured as both the prohormone thyroxine (T4, the form of TH primarily produced by the thyroid follicles, see Fig 6D) and triiodothyronine (T3, the more biologically active form that binds to nuclear TH receptors; Regards et al., 1978; Kikuyama et al., 1993; Shi, 2000; Brown and Cai, 2007). We wanted to test whether the physiological changes stimulated by TH in zebrafish corresponded with periods of elevated TH production or changes in the ratio of T3/T4. We measured T4 and T3 levels at seven stages from larval through juvenile development, staging groups of fish according to the Standardized Standard Length staging criteria (SSL; see Parichy et al., 2009). The relative levels of T3 and T4 remained relatively consistent from one stage to another, except as fish transitioned from 6 to 6.4 SSL. During this period, when anal and dorsal rays first appear (Parichy et al., 2009), we detected a drop in T4 corresponding with an increase in T3 titer (Fig 6A-C). We conclude that there is a shift in TH bioavailability during this mid-larval period, which we have highlighted with grey bars in the graphs of Figs 2, 3, 5–7. Following this shift, T3 availability remained elevated through juvenile development (Fig 6B).

**Figure 6.**
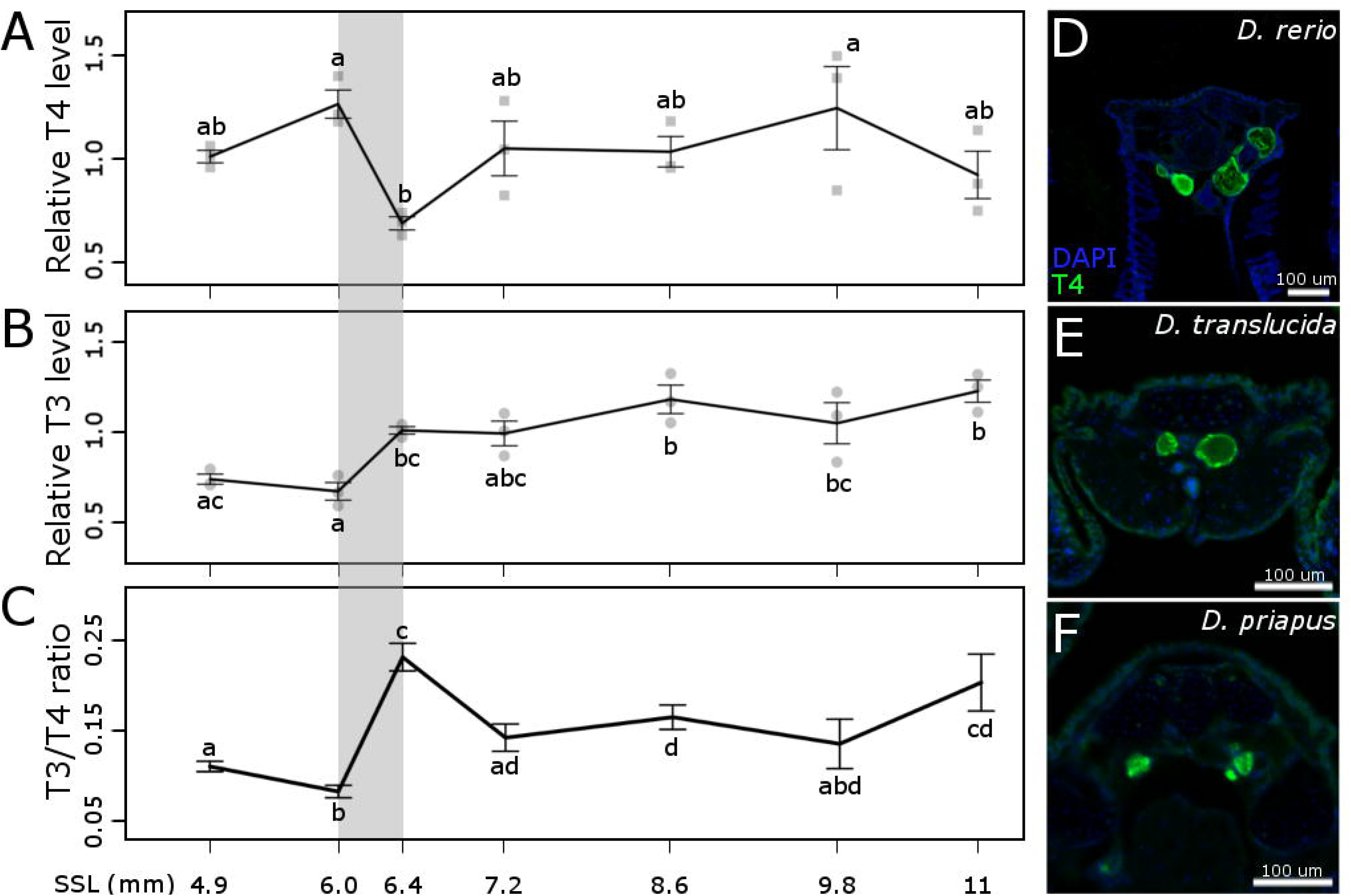
Relative thyroid hormone content during larval development in zebrafish. **A**, T4, thyroxine; **B**, T3, triiodothyronine; **C**, ratio of T3 to T4. All concentrations standardized to body weight. Whiskers represents standard error at each stage. Stages that are not statistically different (as determined by Tukey’s Honest Significant Differences Test, p value cut-off was set at 0.05) are represented with the same letter. For comparison with other figures, grey stripe shows period from 6-6.4 SSL, during which TH availability is elevated. **D-F:** Antibody staining of T4 in zebrafish*J D. priapus* and *D. translucida* adults showing functional thyroid follicles. Images show 12 uM transverse sections of the thyroid region.

**Figure 7.**
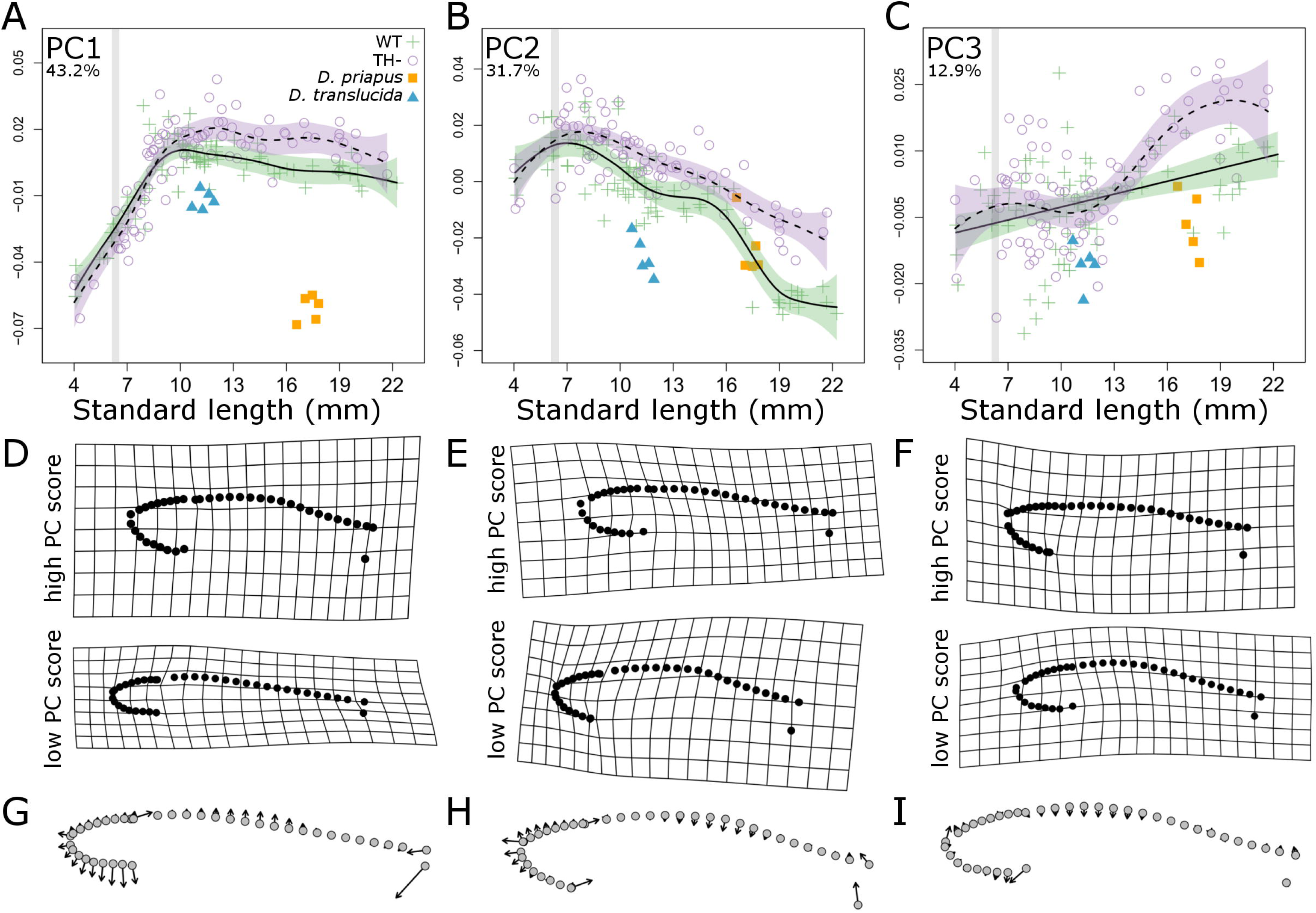
*Danionella* adults resemble larval zebrafish but not TH-zebrafish. **A-C**, Principle component (PC) scores of body shape plotted against standard length for both WT (green cross) and TH-(purple open circle) zebrafish, compared to *D. priapus* (orange square) and *D. translucida* (blue triangle) adults. Percent variation explained by each PC axis is shown on the upper left corner. Developmental trajectory is shown in solid line with purple standard error for WT zebrafish and dashed line with green standard error for TH-zebrafish. For comparison between graphs, grey stripes highlight period from 6-6.4 SL, during which TH availability is elevated. **D-F**, deformation grids showing extreme shapes along each PC. **G-I**, vector plots showing shape change along each PC axis, with arrows pointing from negative to positive PC values.

A previous analysis identified a similar shift in T3/T4 ratio at 10-14 dpf, attributed to higher deiodinase activity (Chang et al., 2012). Indeed, while expression of *thyroglobulin* remains relatively stable from 10-20 dpf, *deiodinase 2* expression increases two-fold (Vergauwen et al., 2018). However, these patterns are difficult to perfectly correlate with our measurements, as dpf (used in these previous analyses) is an unreliable proxy for development (Parichy et al., 2009; McMenamin et al., 2016).

Shifts in allometric trajectories of WT fish occur at 8-10 mm SL (Fig 2A, B; Fig 3C, D). In contrast, the mid-larval shift in T3/T4 ratio occurs at a considerably smaller size (6.4 SSL) and earlier developmental stage (Fig 6), so it seems unlikely that the change in TH ratio directly stimulates the shifts. Nonetheless, the earliest differences between TH- and WT fish are detectable only after ~8 mm SL in allometric relationships (Fig 3B, D) and neuromast numbers (Fig 5B, E). Rather than stimulating a discrete period of profound physiological change (like in metamorphic amphibians), ongoing exposure to TH during late larval periods appears to gradually shift developmental trajectories. TH- zebrafish maintain inappropriately longer periods of larval growth patterns; rather than maintaining larval shapes, TH- zebrafish ultimately achieve novel adult phenotypes.

### Adult *Danionella* resemble larval zebrafish

*Danionella* maintain numerous characteristics of larval zebrafish, and are described as paedomorphic (Britz et al., 2009; Britz and Conway, 2016). We tested the prediction that *Danionella* body shape should more closely resemble larval rather than adult zebrafish, by reanalyzing our morphometric datasets to include *Danionella translucida* and *D. priapus. Danionella* adults of both species exhibited lower PC1 scores than similarly-sized zebrafish (blue triangles and orange squares, Fig 7), showing immature body shape relative to similarly-sized zebrafish. Both species of *Danionella* also showed proportionally smaller heads and eyes for their body length (blue triangles and orange squares, Fig 2). Thus, the two *Danionella* species indeed maintain certain morphologies similar to those of larval zebrafish, and are likely to have more isometric patterns of growth during development.

Although *D. translucida* were smaller than *D. priapus*, the two species had similar total numbers of neuromasts (~300; Fig 5C). The size of the *D. priapus* neuromast population was comparable to that of similarly-sized WT zebrafish, while *D. translucida* had more total neuromasts, with a greater relative density in the trunk (Fig 5B-C, E).

### *Danionella* do not resemble hypothyroid zebrafish, and possess functional thyroid follicles

Since TH is involved in the most well-known examples of paedomorphosis (Gould, 1977; Johnson and Voss, 2013; De Groef et al., 2018), we tested the possibility that TH metabolism was disrupted in *Danionella* species. First, we asked if *Danionella* appeared more similar to TH-than to WT zebrafish. In terms of morphometrics (Fig 7) and ratios of head and eye to SL (Fig 3), both *Danionella* species actually more closely resembled similarly-sized WT than TH- zebrafish. Similar to TH- zebrafish, the two *Danionella* species did not form elongated canal neuromasts (Fig 4D, H). However, *Danionella* show fewer cranial neuromasts and more trunk neuromasts than similarly-sized WT zebrafish, in contrast to the patterns observed in TH- zebrafish (Fig 5). In all, the two *Danionella* species did not resemble TH- zebrafish. To further confirm that hypothyroidism did not underlie developmental truncation in these species, we examined the thyroid follicles, which were intact and showed T4 production comparable to that in zebrafish (Fig 6D-F).

## Conclusions

We identified shifts in allometric trajectories as WT zebrafish transition from larva to juvenile, and some aspects of these shifts showed dependence on TH. However, unlike hypothyroid amphibians, which maintain larval morphology through adulthood, TH- zebrafish inappropriately maintain larval growth trajectories and ultimately achieve novel adult phenotypes. We conclude that TH normally coordinates multiple aspects of the transition from larval to juvenile developmental programs in zebrafish. We interpret these hormonally-mediated changes to constitute a protracted, subtle metamorphosis, in contrast to the sudden and profound metamorphosis of ecologically multiphasic species. Based on its dependence on thyroid hormone, we posit that this subtle metamorphosis is indeed evolutionarily homologous to the profound metamorphosis of ecologically multiphasic vertebrates.

Consistent with the idea that *Danionella* are paedomorphic, two *Danionella* species analyzed showed several similarities to larval WT zebrafish in their body proportions and their lateral line. However, *Danionella* bore little resemblance to TH- zebrafish, and we found that they produced T4. We conclude that, unlike in paedomorphic amphibians, disruptions to the thyroid axis do not underlie paedomorphosis in these species.

## EXPERIMENTAL PROCEDURES

### Fish rearing

Zebrafish were of the line *Tg(tg:nVenus-v2a-nfnB)* (McMenamin et al. 2014), which allows conditional ablation of the thyroid. Transgenic fish were treated 4-5 days post fertilization (dpf) with DMSO (for control euthyroid fish, WT) or DMSO + metronidazole (to ablate the thyroid follicles and produce hypothyroid fish, TH-), following McMenamin et al (2014). All fish were reared at 28°C with a 12:12 light:dark cycle and fed three times per day with marine rotifers, *Artemia* and pure *Spirulina* flakes (Pentair). This diet was chosen to minimize introduction of exogenous TH. *Danionella translucida* was a gift from Adam Douglass, University of Utah; *D. priapus* was purchased through The Wet Spot (Portland OR). Both species of *Danionella* were fed rotifers, *Artemia* and Gemma micro (Skretting).

### Live staining of neuromasts

The fluorescent mitochondrial stain 4-(4-Diethylaminostyryl)-1-methylpyridinium iodide (DiASP, AdipoGen) was used to visualize neuromasts (Collazo et al., 1994). Individual fish were incubated in 0.0024% DiASP in fish system water with 0.02% Tricane (Western Chemical) for 5 minutes, then transferred to 2% methylcellulose and imaged with an Olympus DP74 camera mounted on an Olympus SZX16 stereoscope with a FITC filter. Standard length (SL) was measured using ImageJ (Schneider et al., 2012). Neuromasts were counted manually from digital images; all counts were performed by the same individual to reduce variability.

### Scanning Electron Microscopy

WT and TH- adult zebrafish were fixed in 4% paraformaldehyde (Sigma) overnight at 4°C, rinsed several times with PBS, and dehydrated in an ascending series of ethanol. Samples were critical point dried in 100% ethanol, coated with Au-Pd and imaged with a JEOL JCM-6000PLUS NeoScope Benchtop SEM.

### Fixation and skeletal staining

Specimens were euthanized by an overdose of Tricane, then fixed in 4% paraformaldehyde (Sigma) overnight at 4°C. Specimens were rinsed several times with PBS and dehydrated in 70% ethanol. Following (Walker and Kimmel, 2007), skeletons were stained overnight in 0.5% Alizarin red (Sigma) and 0.02% Alcian Blue (Sigma) with 60 mM MgCl2 in 70% EtOH. Stained specimens were bleached with 3% hydrogen peroxide in 1% potassium hydroxide (KOH), then were cleared by gradually transitioning through an ascending glycerol series from 25% to 80%. Prior to imaging, cleared specimens were incubated in PBS overnight, and then transferred into 2% methylcellulose for positioning and imaged as above for morphometric analysis.

### Morphometrics

Homologous anatomical landmarks and curves that outline the body were placed manually (by a single individual) on lateral images of fish using the R package Stereomorph (Olsen and Westneat, 2015). A list of the specific landmarks can be found in the supplementary materials (supplementary Table S1). Landmark data was then analyzed with standard geometric morphometrics methods using the R package Geomorph (Adams and Otárola-Castillo, 2013).

### Head size and eye size

Head size was measured in images of cleared and stained specimens in lateral view. Head size was measured as the distance from the tip of the snout/rostral edge of premaxilla to the posterior tip of the occipital bone. Two eye measurements, the dorsal-ventral height, and the anterior-posterior length were obtained, and then averaged as the eye-size measure. Measurements were performed by a single individual to reduce variability.

### Thyroid staining

To image thyroid follicles, adult zebrafish, *D. priapus* and *D. translucida* were fixed in 4% paraformaldehyde (Sigma), embedded in O.C.T. Compound (Fisher Scientific), and sectioned on to 12 µm. Slides were washed with 0.1% Tween in PBS (PBSt), blocked with 10% heat-inactivated goat serum in PBSt, then incubated at 4°C overnight with a rabbit T4 antibody (1:1000, Fitzgerald). Slides were then washed, incubated with a secondary antibody (goat anti-rabbit FITC diluted 1:1000, Fitzgerald), stained with DAPI (1ug/ml) for 20 minutes, washed and imaged on an Olympus IX83 microscope equipped with a Hamamatsu ORCA Flash 4.0LT camera.

### Quantification of thyroxine (T4) and triiodothyronine (T3)

Live WT zebrafish were anesthetized and staged following the adult normal table (Parichy et al. 2009). Seven stages were chosen to capture major processes during the entire transition from larva to juvenile (see Supplemental Table 2), and fish were staged by the SSL convention (Parichy et al., 2009). Samples were prepared following McMenamin et al. (2014); briefly, staged fish were pooled, euthanized by an overdose of tricaine, blotted dry, and weighed in microcentrifuge tubes. Tissue was suspended in cold methanol with 1 mM 6-propyl-2-thiouracil (PTU; Sigma) and homogenized by sonication; homogenized samples were stored at −80°C until analysis. Samples were centrifuged, re-extracted with additional PTU-containing methanol. Supernatants were pooled and dried with a CentriVap Concentrator (Labconco) for ~5 hours, then resuspended in PBS. Samples were run on T4 and T3 ELISA kits (Diagnostic Automation Inc.) with two technical replicates and with three biological replicates for each stage. Plates were analyzed on a PerkinElmer Victor X5; mean absorbance value from three readings was used for subsequent analysis. Standard curves were generated with 4-parameter logistic regression via online analysis software (www.elisaanalysis.com).

### Statistics

All statistical analyses were performed in in R (version 3.4.1). Differences in thyroid hormone levels along ontogeny were first examined using ANOVA, followed by Tukey’s Honest Significant Distance Test. Developmental trajectory was estimated with the spm function in the package SemiPar using default settings (Carroll et al., 2003). Developmental trajectories were considered to be distinct when the ranges of standard error did not overlap between the groups. For example, in Fig 2A, the standard error ranges always overlap (the green and purple bands overlap) and the models are not considered different. In Fig 2B, the standard error ranges of two models diverges at ~9 mm SL.

## ACKNOWEDGEMENTS

Thank you for discussion and technical support from Catherine May, Angela Mauri Stacy Nguyen, Pranav Parikh, Mary Xu and all the members of the McMenamin Lab. Thank you to Jacqueline Webb and another anonymous reviewer for their insightful comments. Thank you to Adam Douglass, David Parichy, the Boston College Statistics Department and the Boston College Animal Care staff. We gratefully acknowledge support from NIH R00GM105874, R03HD091634, Burroughs Wellcome Grant 1017439 and NSF CAREER 1845513.

---

**Supplementary Table S1.**
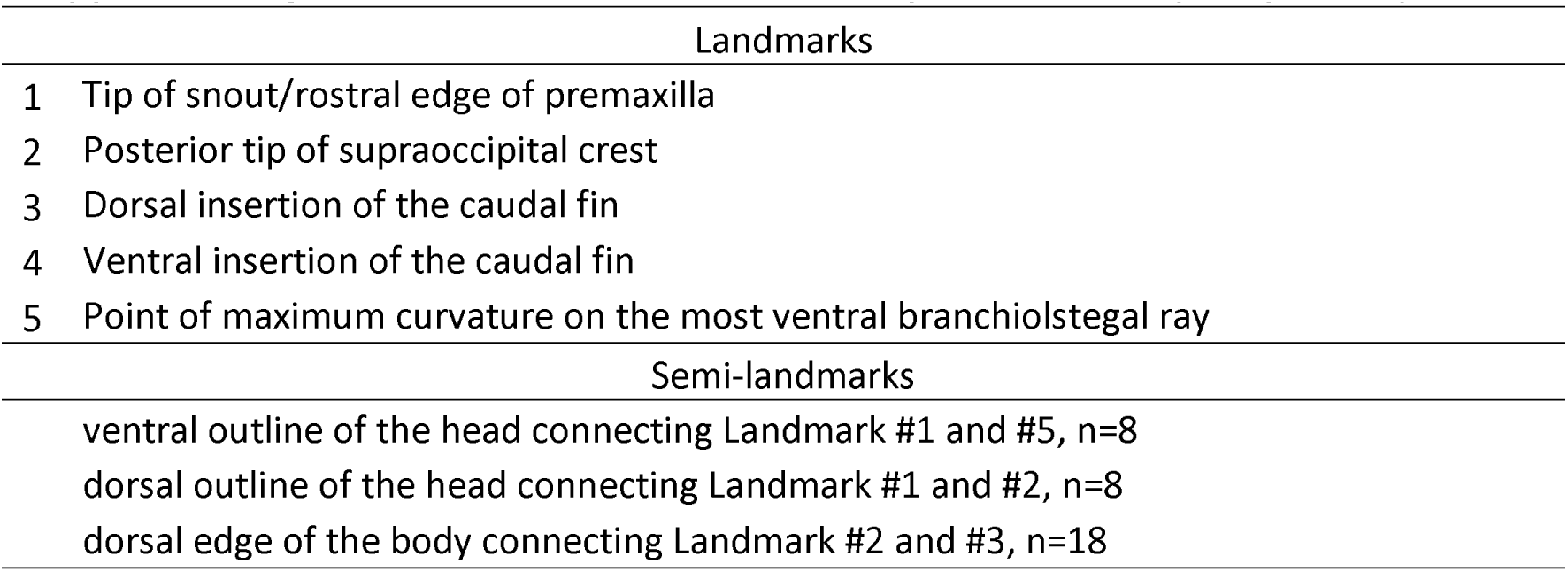
Landmarks used in morphometric body shape analysis.

**Supplementary Table S2.** Summary of fish samples used for ELISA.

| SSL | Stage | SL (mm) | Age (± SD) | n | average weight (g) per fish |
| --- | --- | --- | --- | --- | --- |
| 4.9 | CR | 4.9-5.1 | 12 ± 2.9 | 1318 | 0.000893649 |
| 6 | aSB | 5.9-6.2 | $13.5 \pm 2.5$ | 654 | 0.001709021 |
| 6.4 | DR | 6.4-6.6 | $15 \pm 2.5$ | 403 | 0.003138213 |
| 7.2 | PB | 7.2-7.5 | $16 \pm 2.7$ | 294 | 0.004842177 |
| 8.6 | PR | 8.5-8.7 | $20 \pm 1.3$ | 149 | 0.008157047 |
| 9.8 | SP | 9.5-9.6 | $26.5 \pm 2.1$ | 98 | 0.013067347 |
| 11 | J | 10.9-11.7 | $32 \pm 0.5$ | 58 | 0.022691379 |

